# A mechanistic model of the neural entropy increase elicited by psychedelic drugs

**DOI:** 10.1101/2020.05.13.093732

**Authors:** Rubén Herzog, Pedro A.M. Mediano, Fernando E. Rosas, Robin Carhart-Harris, Yonatan Sanz Perl, Enzo Tagliazucchi, Rodrigo Cofre

**Affiliations:** Centro Interdisciplinario de Neurociencia de Valparaíso, Universidad de Valparaíso, Pje Harrington 287, 2360103 Valparaíso, Chile; Department of Psychology, University of Cambridge, Cambridge CB2 3EB, UK; Centre for Psychedelic Research, Department of Brain Science, Imperial College London, London SW7 2DD, UK; Data Science Institute, Imperial College London, London SW7 2AZ, UK; Centre for Complexity Science, Imperial College London, London SW7 2AZ, UK; Buenos Aires Physics Institute and Physics Department, University of Buenos Aires, Buenos Aires, Argentina; National Scientific and Technical Research Council, Buenos Aires, Argentina; Universidad de San Andres, Buenos Aires, Argentina; CIMFAV-Ingemat, Facultad de Ingeniería, Universidad de Valparaíso, Valparaíso, Chile

## Abstract

Psychedelic drugs, including lysergic acid diethylamide (LSD) and other agonists of the serotonin 2A receptor (5HT2A-R), induce drastic changes in subjective experience, and provide a unique opportunity to study the neurobiological basis of consciousness. One of the most notable neurophysiological signatures of psychedelics, increased entropy in spontaneous neural activity, is thought to be of relevance to the psychedelic experience, encoding both acute alterations in consciousness and mediating long-term effects. However, no clear mechanistic explanation for this ‘entropic’ phenomenon has been put forward so far. We sought to do this here by building upon a recent whole-brain model of serotonergic neuromodulation, to study the entropic effects of 5HT2A-R activation. Our results reproduce the overall entropy increase observed in previous experiments *in vivo*, providing the first model-based explanation for this phenomenon. We also found that entropy changes were not uniform across the brain: entropy increased in some regions and decreased in others, suggesting a topographical reconfiguration mediated by 5HT2A-R activation. Interestingly, at the whole-brain level, this reconfiguration was not well explained by 5HT2A-R density, but related closely to the topological properties of the brain’s anatomical connectivity. These results help us understand the mechanisms underlying the psychedelic state and, more generally, the pharmacological modulation of whole-brain activity.

## Introduction

Psychedelic drugs provide a privileged opportunity to study the mind-brain relationship, and promise to revolutionise some of our current mental health treatments.^1–3^ However, while some aspects of the neurochemical action of psychedelics at the neuronal and sub-neuronal level are well known,^4, 5^ our current understanding of their action at the whole-brain level, is still very limited. A deeper understanding of the mechanisms that trigger the changes in conscious experience produced by psychedelics would greatly advance our knowledge of human consciousness, and medical development of psychedelics.

At a subjective level, serotonergic psychedelics (including LSD, dimethyltryptamine [DMT] and psilocybin, among others) potentially modulate mood, cognition, perception and self-awareness. At a neurophysiological level, recent research has identified (among many) two particularly prominent signatures of the psychedelic state: an overall disregulation of neural population activity, most clearly seen as suppression of spectral power in the alpha (8-12 Hz) band;^6–8^ and an increase in the signal diversity of the neural activity, measured through the information-theoretic notion of entropy.^9^ In particular, this acute entropy increase has been linked to both short- and long-term effects of the psychedelic experience, including certain aspects of the reported subjective experience^9^ and subsequent personality changes.^10^.

Interestingly, the opposite effects have been reported for states of loss of consciousness, where a strong *decrease* in brain entropy has been repeatedly observed. This seems to be a core feature of loss of consciousness, generalising across states such as deep sleep,^11^ general anaesthesia,^12^ and disorders of consciousness.^13^ Together, these divergences are yielding converging evidence that entropy and related measures offer simple and powerful indices of conscious states.

Relatedly, Carhart-Harris and colleagues have put forward the *Entropic Brain Hypothesis* (EBH) an entropy based theory of conscious states.^1, 14^ The EBH proposes the simple, yet powerful idea that the variety of states of consciousness can be indexed by the entropy of key parameters of brain activity, or in other words, that the richness of subjective experience is directly related to the richness of on-going neural activity, where richness can be defined most simply as diversity. Investigating the neurobiological origins of such changes in brain activity is therefore a key step in the study of altered states of consciousness.

Multiple experiments in humans and animal models have established that the mind-altering effects of psychedelics depend on agonism specifically at the serotonin 2A receptor (5HT2A-R).^15–17^ A recent simulation study involving whole-brain computational modelling confirmed that the topographic distribution of 5HT2A-R in the human brain is critical to reproduce the functional connectivity dynamics of human fMRI data recorded under the acute effects of LSD.^18^ Here we build upon this model to characterise the interplay between the entropy of brain signals, the distribution of 5HT2A receptors, and structural connectivity properties of the brain, with the overarching goal of explaining the sources of entropic effects, and thus altered consciousness, elicited by psychedelic drugs.

## Results

We simulated whole-brain activity using the Dynamic Mean-Field (DMF) model by Deco *et al.*,^19^ using parameter values fit to reproduce the dynamics of fMRI recordings in humans during wakeful rest, as well as under the acute effects of LSD.^18^ The model consists of interacting pools of excitatory and inhibitory neural populations, coupled via long-range excitatory connections informed by the anatomical connectivity of the brain. The DMF model combines a theoretical model of neural and synaptic dynamics with two empirical sources of information about the human brain: the human connectome, i.e. DTI-estimated connectivity between the 90 regions of the AAL^20^ atlas; and average 5HT2A-R expression across the human brain obtained with Positron Emission Tomography (PET) scans^21^.

Each simulation of the DMF model generates 90 time series of excitatory firing rates, one for each region of the AAL atlas (Fig. 1A). These excitatory firing rates have a non-linear dependency on the local excitatory inputs, determined by a frequency-current (*F-I*) curve; and 5HT2A-R activation is modelled as a response-gain modulation of this F-I curve dependent on the receptor density at a given region (Fig. 1B). We simulated the model in two conditions, with and without 5HT2A-R activation, which, in an analogy with neuroscientific terminology, we refer to as placebo (PLA) and 5HT2A-R conditions, respectively. In this way, we obtain 90 time series for each condition, which we keep for further analysis.

**Figure 1.**
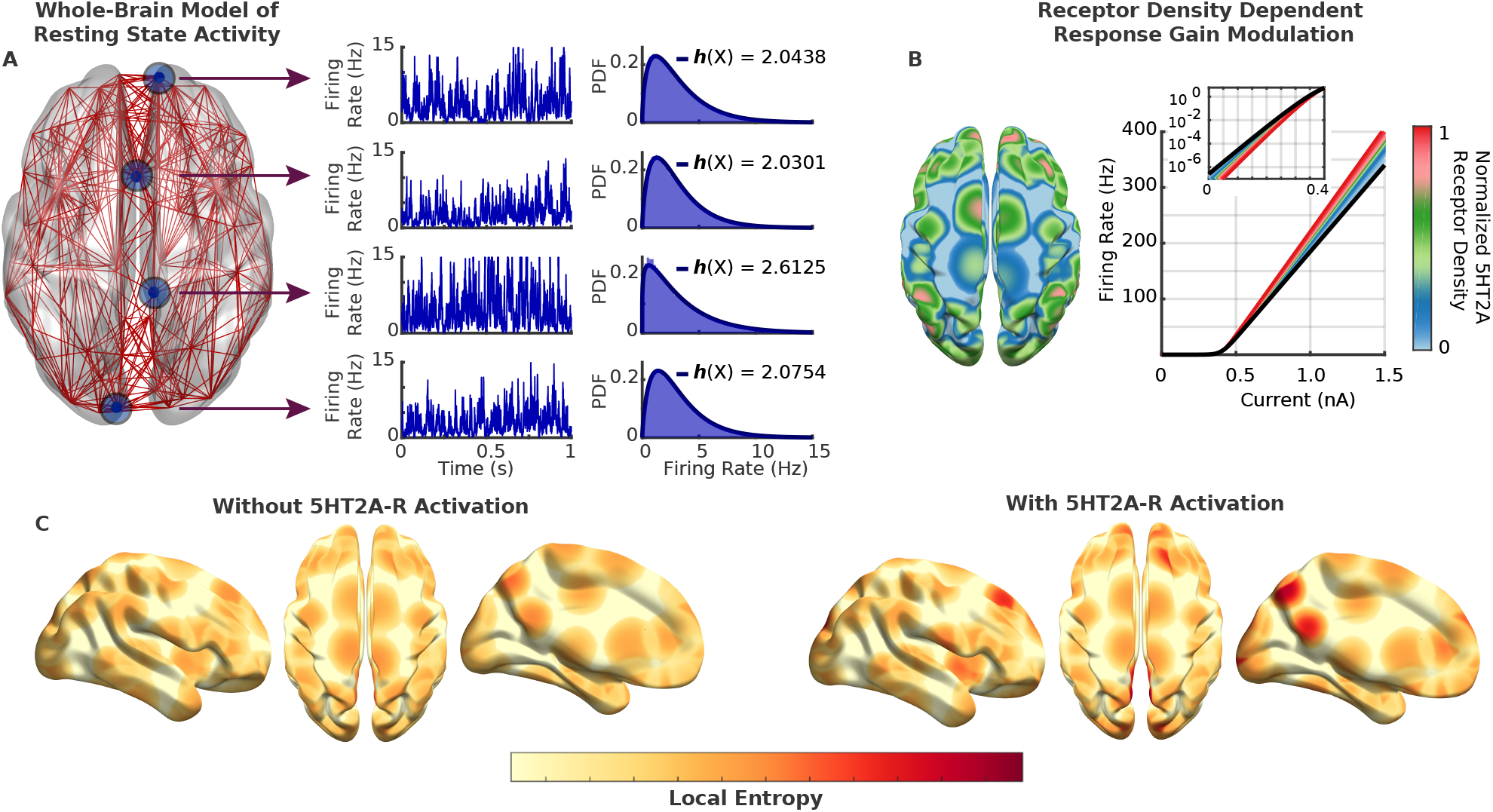
Modelling the effect of 5HT2A-R activation on the whole-brain topographical distribution of entropy. **(A)** Resting state activity is simulated using the Dynamic Mean-Field (DMF) model, in which each region’s activity is represented by a time series of excitatory firing rates. The differential entropy of each region is then estimated, obtaining a topographical distribution of entropy values. **(B)** 5HT2A-R agonism is modelled as a receptor-density-dependent response gain modulation. Black line is the frequency-current (F-I) curve of a population without 5HT2A-R agonism, and coloured curves show the resulting F-I curves of regions with increasing 5HT2A-R agonism. **(C)** 5HT2A-R activation changes the topographical distribution of entropy with respect to resting state activity, which constitutes the main subject of analysis in this study.

Finally, using these time series, we estimate Shannon’s differential entropy for each region under both conditions, yielding a topographical distribution of entropy values (Fig. 1C) that constitutes the main subject of analysis in this study. Further details about model specification and entropy estimation, as well as other methodological caveats, are presented in the Methods section at the end of this article.

### 5HT2A-R activation causes a heterogeneous, non-linear increase in the entropy of simulated brain activity

In our first analysis, we used the DMF model to test the main prediction of the EBH: that 5HT2A-R activation causes an increase in the overall entropy of neural signals (Fig. 2). In line with previous experimental findings with psychedelic drugs,^9^ the model shows a significant increase in the brain’s entropy *h* due to 5HT2A-R activation, with an average entropy of *h*^PLA^ = 2.20 nat in the placebo condition, and *h*^5HT2A^ = 2.30 nat in the 5HT2A-R condition (dashed vertical lines in Fig. 2B, Wilcoxon signed-rank test *p* < 10^−6^, Cohen’s *d* = 0.262 ± 0.011). A closer look at the distribution of entropy changes, however, reveals a more nuanced picture, with some regions increasing and some decreasing their entropy as a result of 5HT2A-R activation (Fig. 2D). This suggests that, according to the model, 5HT2A-R agonism might trigger a complex reconfiguration of the topographically specifc distribution of entropy, and not a mere homogeneous overall increase.

**Figure 2.**
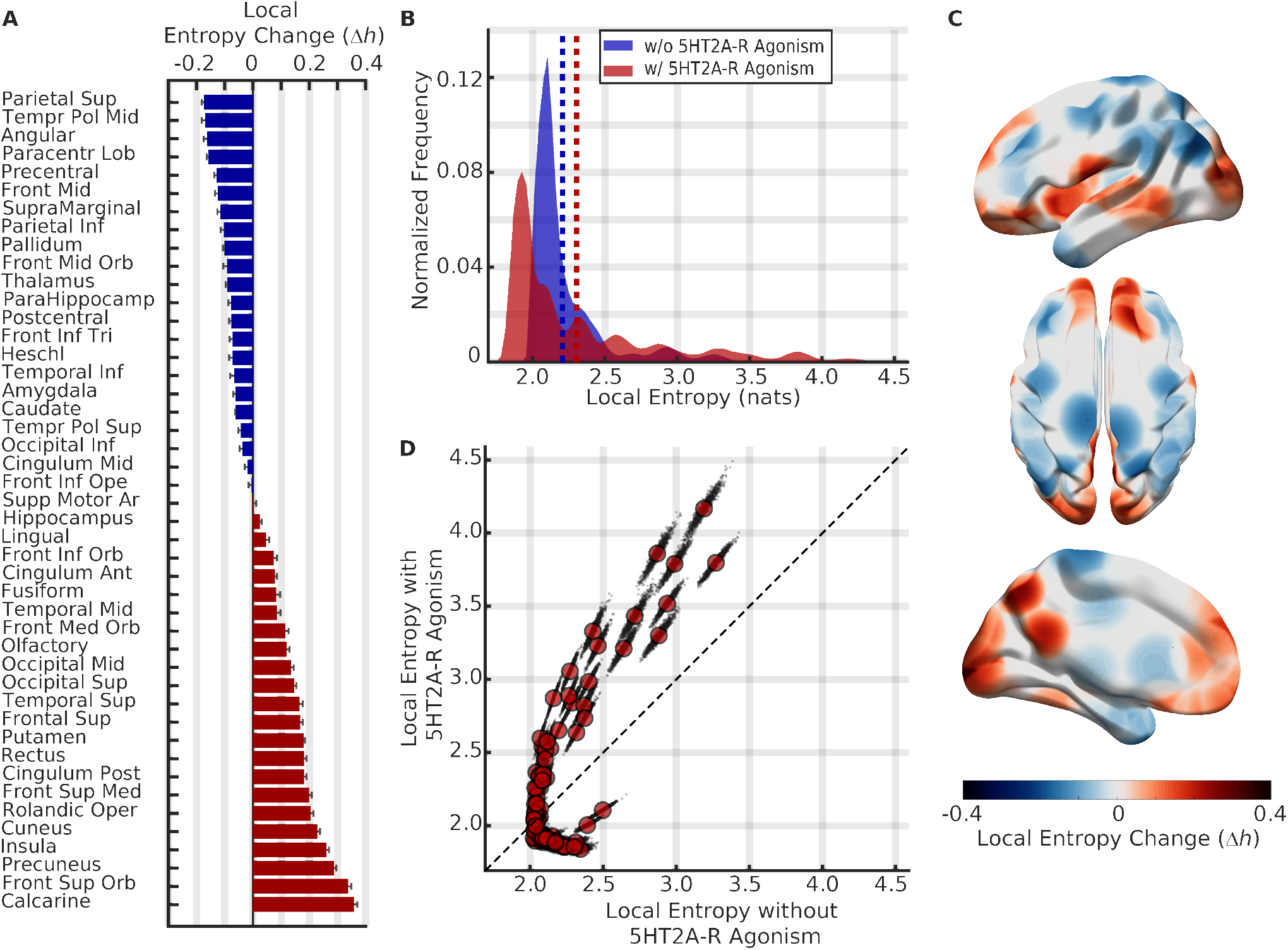
Non-linear heterogeneous increase of entropy following 5HT2A-R activation. **(A)** Effect of 5HT2A-R agonism on the local entropy each of region in the AAL atlas. Bars indicate the (bilateral) average relative change in local entropy, Δ*h*_*n*_, and error bars indicate 1 standard deviation across 1000 simulations. **(B)** Histograms of local entropy values for the condition with (red) and without (blue) 5HT2A-R activation. 5HT2A-R activation increased both the average and the spread of the local entropy distribution. **(C)** Topographical map of entropy changes. Brain regions are coloured according to their Δ*h*_*n*_ values. **(D)** 5HT2A-R agonism changed the topographical distribution of entropy in a heavily non-linear manner. Black dots indicate single simulations and red circles indicate the averages across all simulations.

To investigate this reconfiguration, we analysed the local changes in entropy by plotting the entropy of the *n*^th^ region in both PLA and 5HT2A conditions, denoted by 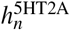 and 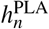 respectively (Fig. 2D). Our results show that 5HT2A-R activation affects local entropy in a highly non-linear manner, especially in regions with low baseline resting state entropy. In particular, in such regions the effect of 5HT2A-R activation could either increase or decrease entropy, such that the local entropy in the 5HT2A-R condition could not always be determined by the region’s baseline entropy. This finding hints towards a more general theme: that local dynamical properties alone are not able to explain the local changes in activity induced by 5HT2A-R agonists like psychedelic drugs. We will explore this phenomenon in depth in the following sections.

Next, we studied the effect of 5HT2A-R agonism on local entropy by considering the relative change scores,

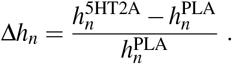

Based on recent *in vivo* experiments with serotonergic psychedelics we expect to find localized entropy increases on occipital, cingulate and parietal regions^9, 10^ as well as several changes on regions belonging to Resting State Networks (RSNs)^7, 22, 23^. Then, we split brain regions by standard anatomical and functional groupings, finding that occipital and cingulate areas tend to show a strong relative increase in their entropy, while parietal and subcortical areas tend to show a decrease (Fig. 2A, Supp. Fig. 1). Additionally, regions participating in the primary visual and Default Mode (DMN) RSNs - including occipital and cingulate areas, respectively - showed a marked tendency to increase their entropy, while regions participating in the Frontoparietal Executive Network (FPN) showed the opposite behaviour. Further discussion about the relation between these results and recent *in vivo* experiments with psychedelic drugs is presented in the Discussion section.

### The topography of entropy changes is explained by local connectivity strength and 5HT2A-R density

Given the results above, and given the potential clinical and neuroscientific relevance of entropy reconfiguration in the psychedelic state, our next task is to elucidate what neurobiological factors underlie such phenomenon. In this section we investigate which structural and dynamical features of the model are able to predict local entropy changes due to 5HT2A-R activation, and what we can learn about the large-scale action mechanism of psychedelic drugs and other 5HT2A-R agonists.

To explain the effects of 5HT2A-R activation, the first natural step is to factor in the density of 5HT2A-R at each region. Somewhat surprisingly, at a whole-brain level, receptor density was a very poor predictor of the entropy change due to 5HT2A-R activation (Fig. 3A, *R*^2^ = 0.078 ± 0.004). In contrast, we found that the connectivity strength (based on the DTI human connectome), defined as the sum of all the weighted links connecting a given region, exhibits a strong correlation with local entropy changes (Fig. 3B, *R*^2^ = 0.801 ± 0.006).

**Figure 3.**
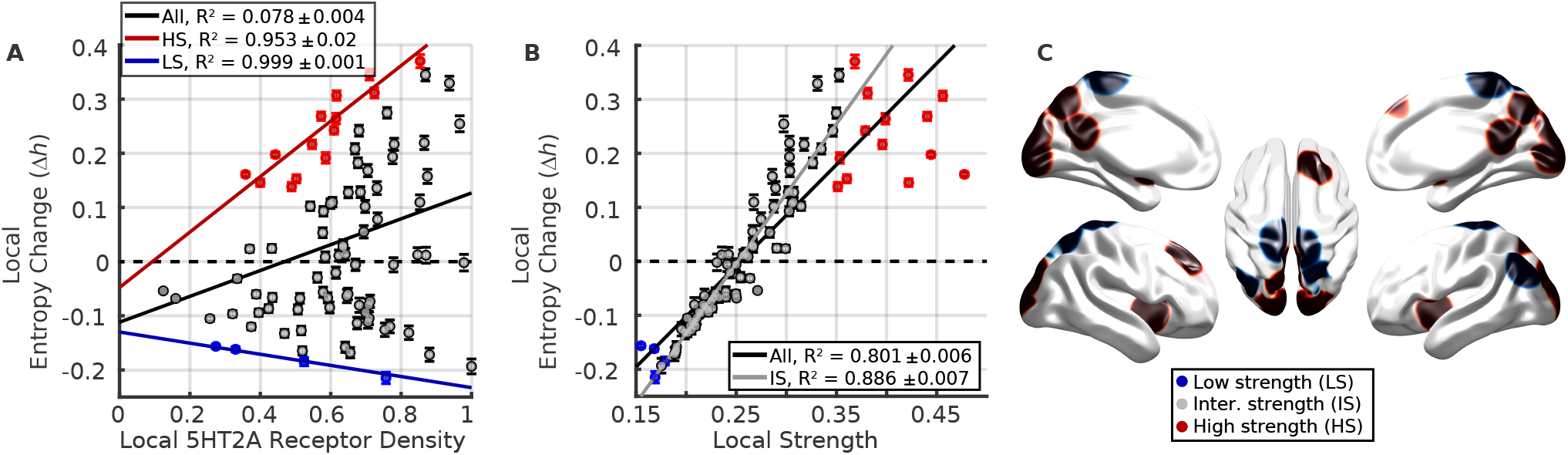
Changes in local entropy are explained best by connectivity strength, then receptor density. **(A)** Changes in entropy were overall independent from receptor density, although **(B)** they were well predicted by the connectivity strength of each region. We split into groups of low (blue), intermediate (grey) and high (red) connectivity strength, only those regions with low and high connectivity strength were well predicted by their receptor density. Regions with intermediate strength show no significant relationship with receptor density, but are even more accurately predicted by their strength. **(C)** Topographical localisation of the three groups, following the same colour code. Low-strength regions are mainly located in the parietal area, while high-strength ones are in occipital and cingulate areas.

By visual inspection, however, it is clear that the relationship between DTI-informed connectivity strength and Δ*h*_*n*_ is linear for intermediate values of strength, but it departs from linearity in both extremes of the distribution. To investigate this phenomenon, we implemented a simple optimisation algorithm to find the set of regions with the strongest linear relationship between strength and Δ*h*_*n*_. This yielded a set of regions with a higher strength-Δ*h*_*n*_ correlation (*R*^2^ = 0.886 ± 0.007, as opposed to *R*^2^ = 0.801 ± 0.006 for the whole-brain), and induced a partition of all brain regions into three groups: those with intermediate strength (IS, 76 ± 4 brain regions, grey dots in Fig. 3A), high strength (HS, 12 ± 3 regions, red dots), and low strength (LS, 2 ± 2 regions, blue dots).

Building up on this partition, we studied the effect of receptor density on those regions where entropy change could not be predicted from connectivity strength - the HS and LS groups. We found a strong relationship between receptor density and entropy changes for both the HS and LS groups, which resulted in positive (*R*^2^ = 0.95 ± 0.02) and negative (*R*^2^ = 0.999 ± 0.001) correlation, respectively. On the contrary, receptor density does not predict entropy changes for areas in the IS category. This shows a complementary role of density and strength in LS, IS, and HS regions: entropy changes in HS and LS regions strongly depend on the receptor density, but not connectivity strength; while changes in IS regions depend on connectivity strength, but not receptor density.

Overall, to assess the predictive power of connectivity strength and the receptor density on local entropy changes, we built a linear mixed model for Δ*h*_*n*_ using connectivity strength, 5HT2A-R density, and the aforementioned three-way separation of brain regions as the predictor variables. Together these variables explain 95.91 ± 0.01% of the variance of Δ*h*_*n*_, confirming that they provide an accurate model for predicting the entropic effects of 5HT2A-R activation. This suggests that psychedelic drugs and other 5HT2A-R agonists do not have a simple, localised effect on brain activity, but instead amplify the fundamentally collective, emergent properties of the brain as a complex system of interacting elements.

### The specific connectivity strength distribution explains relative changes in entropy

As a final analysis, we set out to investigate exactly which topological properties of the brain’s structural connectivity explain the observed changes in entropy. With this exercise, we aim to answer two questions: whether any property simpler than connectivity strength *can* explain the results; and whether any property more complicated than connectivity strength *is needed* to do so.

To this end, we ran further simulations of the DMF model using suitable null network models of the human connectome (Fig. 4A), with the 5HT2A-R density map held fixed. In particular, we used three null models designed to test increasingly strict null hypotheses that preserved different network attributes of the original connectome: i) the overall density and strength (RAND, Fig. 4B), ii) the degree distribution (degree-preserving randomisation [DPR], Fig. 4C); and iii) the strength distribution (strength-preserving randomisation, [DSPR], Fig. 4D). We simulated the DMF model in these surrogate networks, with and without neuromodulation, and computed the resulting entropy changes Δ*h*_*n*_.

**Figure 4.**
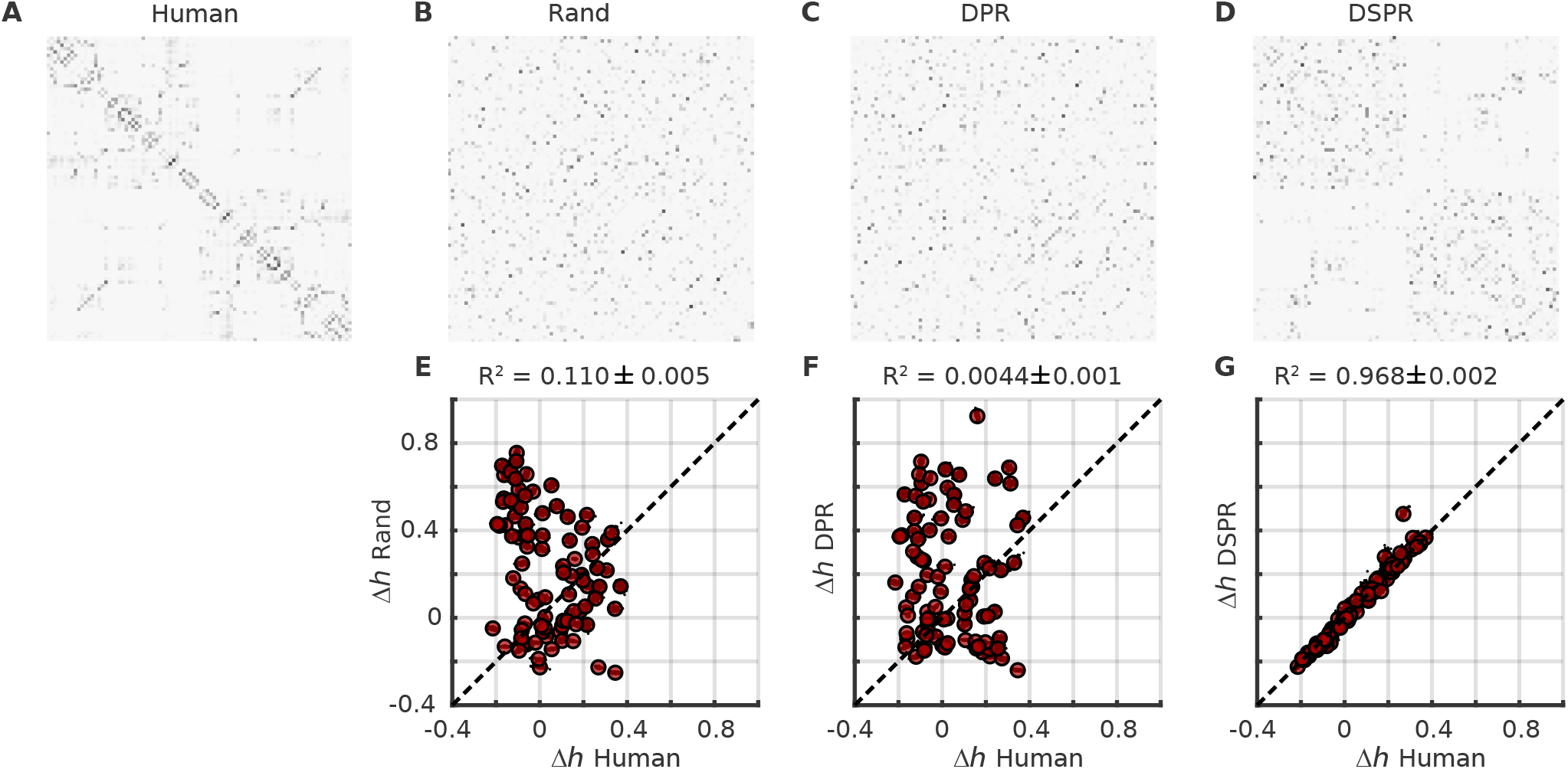
Relative changes in entropy are reproduced by a strength-preserving null model of the connectome. **(A)-(D)** Connectivity matrices used to control the role of local properties of the connectome on Δ*h*_*n*_. See main text for the description of the matrices and randomisation algorithm. **(E)-(G)**. Scatter plots of Δ*h*_*n*_ for the human connectome against the three null models. DSPR yielded almost the same results than the human connectome, showing that only local network properties of human connectome are sufficient to capture the effect of 5HT2A-R activation.

Our first result is that random and degree-preserving surrogate networks are unable to reproduce the entropy changes observed in the human connectome: when compared against the Δ*h*_*n*_ values obtained from the unperturbed network, neither of them produce values close to the original (Figs. 4E, 4F). This result asserts the findings in the previous section, showing that indeed node strength plays an important role in shaping the global pattern of entropy change associated with the action of psychedelics and other 5HT2A-R agonists. Simpler topological features, like the degree distribution, are not enough to explain such changes.

Perhaps more interestingly, the connectivity strength-preserving surrogate networks not only reproduced the entropy changes, but did so almost exactly (Fig. 4G). Furthermore, we repeated the analyses in the previous section trying to predict Δ*h*_*n*_ using other local connectivity measures (e.g. betweenness, eigenvector centrality), and none of them produced a model as accurate as strength (Supp. Fig. 2). Together, these results show that, once the receptor distribution is fixed, the network strength distribution is both *necessary* and *sufficient* to explain the entropic effects of 5HT2A-R activation. Of course, this is not to say that strength explains every aspect of 5HT2A-R action - other topological network features are known to mediate transitions of consciousness in other contexts,^24^ and investigating which network properties explain high-order dynamical signatures^25^ of psychedelics remains an exciting avenue of future work.

## Discussion

In this study we investigated the brain entropy changes induced by serotonergic psychedelics, by simulating whole-brain resting state activity with and without 5HT2A-R activation. In contrast to empirical studies, which usually only have access to coarse-grained fMRI or M/EEG data, our approach allows us to study firing rates of brain regions and hence to directly assess the effect of 5HT2A-R agonism at the level of neural population activity.

In agreement with the Entropic Brain Hypothesis,^14^ the model shows a significant increase of global brain entropy, albeit in a highly inhomogeneous and non-linear reconfiguration. The diversity of effects stresses the importance of extending the scope of the EBH from a simple level-based approach towards a multi-dimensional perspective, which might better characterise the richness of 5HT2A-R activation effect both on the brain and consciousness.

These results have important consequences for our understanding of consciousness, neurodynamics, and the psychedelic state. In the following, we highlight some of those implications, and propose possibilities for future work.

### 5HT2A-R-induced entropy changes are regionally heterogeneous

Our main result in this study is a mechanistic explanation of the main prediction of the EBH that serotonergic psychedelics increase the entropy of brain activity. However, one of the main takeaways is that this overall increase is tied to a spatially heterogeneous reconfiguration, rather than a globally consistent increase, in entropy.

Our simulation results show that 5HT2A-R activation triggers an entropy increase in sensory areas, as observed in fusiform and olfactory cortices, as well as the primary visual RSN and occipital regions (Supp. Fig. 1). This agrees with the increased perceptual ‘bandwidth’ that characterises the psychedelic state,^26^ as higher entropy might be related to richer perceptual experience in a given moment of consciousness - potentially related to reduced gating. As a matter of fact, all the primary visual RSN regions (with the exception of the lingual area) are part of the high-strength group, suggesting that the effect of psychedelics on perceptual experience might be directly related to 5HT2A-R density in those regions. The localisation of the entropy increases may relate to domain specific changes in consciousness, which could be interpreted as consistent with a recent dimensions of consciousness characterization of the psychedelic experience.^27^

### Comparison to *in vivo* experiments with psychedelic drugs

Throughout the paper, we have focused on one particular signature of 5HT2A-R agonists on the brain - a global increase in average entropy. But much more is known about the effect of psychedelics on the brain, and studying these more nuanced effects is key to understanding the rich phenomenology of the psychedelic state. In this section we deepen this connection by providing a more detailed comparison between the behaviour of the model and experimental results with psychedelic drugs.

Our findings are in agreement with earlier studies where the effect of psychedelic drugs on the topographical distribution of entropy-related measures was correlated with subjective effects. For example, Schartner *et al.*^9^ studied the effect of LSD, psilocybin, and ketamine on the entropy rate of binarized MEG signals, and found localised increases in entropy rate on occipital and parietal regions. Similarly, Lebedev *et al.*^10^ analysed the sample entropy of fMRI recordings of subjects under the effect of LSD, finding localized increases on frontoparietal, medial occipital, posterior and dorsal cingulate regions. Many of those regions showed a consistent increase of their entropy with 5HT2A-R agonism in our study (Fig. 2A).

Moreover, many of those regions actually belong to the high connectivity strength group (c.f. Fig. 3), which suggests that their entropy increase in experimental data might be directly related to the high 5HT2A-R density in those regions. Together, these findings support the conclusion that the DMF model, once optimised, can reproduce not only functional connectivity,^18^ but also some of the most salient localised entropy increases found on *in vivo* human studies up to date.

Also, on a more fundamental level, our findings suggest that psychedelics dysregulate the functional organisation of the brain with an especially focal and pronounced action on highly anatomically connected brain regions. This finding has very important implications for our understanding of global brain function where a possible inference on this would be that the maintenance of regular (i.e. non entropic) activity in such key connector hubs is critical for the maintenance of normal waking consciousness, and moreover, perturbing the activity in these specific regions (via 5HT2A-R agonism) has particularly profound implications for the quality of conscious experience.

On a separate line of inquiry, there is strong evidence associating the DMN to high-level cognitive functions, including the sense of space, time and recognition of (self) boundaries. Disruptions to the DMN have been linked to fundamental changes in experience, such as ego dissolution^14, 28^. Our simulation results show that 5HT2A-R activation increases the entropy of all DMN regions (with the exception of the angular cortex), which is consistent with the reported decrease in the DMN network integrity^29^ induced by psychedelic drugs. In contrast, low-level motor functions such as motor regulation remain largely unaffected during the psychedelic state,^30^ which is consistent with the modest entropy changes observed in the lingual and superior motor areas.

Finally, it is worth noting that both angular regions showed particularly important decreases in local entropy, specially in the left hemisphere (Fig. 2A). Damage to this region is associated with impairments on language processing, and electrical stimulation can induce out-of-body experiences^31^ (both experiences can feature within psychedelic states^32^). Since both structural damage and electrical stimulation can be related to entropy reduction (e.g. by stimulus-driven synchronisation), these experimental findings are consistent with the strong entropy decrease in angular regions predicted by the model. We speculate that the reduced entropy observed on the angular region induced by 5HT2A-R agonism could be related to a smaller local bandwidth, which in turn might impair the multi-modal and integrative functions of this region.^31^

### Current limitations and future research

The approach employed in this work presents certain limitations related to several aspects of the simulation and analysis. Acknowledging and understanding these limitations can help us extend and improve our approach, while introducing new questions in the field of psychedelic computational neuroscience.

To our knowledge, the DMF model is the only whole-brain model that implements neuromodulation and is capable of reproducing neuroimaging data in the placebo and psychedelic states. Nonetheless, it makes some important simplifications that are worth discussing. At the network level, the DTI-based connectome used here is known to be incomplete, thus improvements could be made to the model parameters of brain connectivity.^33^ At the dynamical level, the DMF model models neuronal populations as perfectly homogeneous within a given region, and it is known that finer-grained local structure of certain brain regions, such as the lattice structure of the primary visual cortex, is likely to be key to explaining certain subjective effects of the psychedelic state (e.g. lattice structure in the visual cortex and geometric visual hallucinations^34^). Additionally, the version of the DMF model used here only considers 5HT2A-R agonism, while classic serotonergic psychedelic drugs also have high binding affinities for other receptors (e.g. in the case of LSD, the D1 and D2 dopamine receptors^35^).

These simplifications do not prevent the model from reproducing statistical features of brain signals under the placebo and LSD conditions, but could result in an inability to reproduce finer aspects of the dynamics of the whole-brain activity in these conditions. Extending the model to reproduce other dynamical signatures of psychedelics (like alpha suppression^8^ or reduced directed functional connectivity^36^) constitutes a natural extension of this work.

Another potentially fruitful line of future work involves making more detailed comparisons with *in vivo* psychedelic neuroimaging data, and, potentially, subjective experience reports. For example, one natural option would be to use forward models of fMRI^37^ or M/EEG^38^ to bridge between the firing rates produced by the DMF model and other data modalities, to produce simulated data that is more directly comparable with available empirical data.

Another exciting possibility is to explore model parameters to examine potential non-linearities in their implications for different relevant aspects of brain function. For example, there are some reasons to believe that the dose-response relationship is non-linear for psychedelics and that over a certain threshold dosage (level of 5HT2A-R stimulation) - new subjective and global brain function properties can feature^39^.

Most interestingly, a potentially very useful extension of this work would be to include individual subject-level connectome and receptor data to build personalised models of response to psychedelic drugs. This would enable a much more comprehensive modelling framework, capable of correlating structural brain features with subjective experience reports. Such a framework could potentially make individualised predictions of the action of serotonergic psychedelics on specific individuals, aiding patient stratification and treatment customization.^2^

Finally, it is worth noting that all our analyses here are based on the univariate statistics of individual brain regions, not including any correlation or information flow between them. However, it is known that some high-level subjective effects of psychedelics (such as complex imagery^7^ and ego dissolution^28^) are related to network, as opposed to single region, dynamics. Therefore, building a richer statistical description of the brain’s dynamics using recent information-theoretic tools (such as multivariate extensions of mutual information^25^) remains an exciting open problem.

### Final remarks

In this paper we have provided the first mechanistic explanation of the neural entropy increase elicited by psychedelic drugs, using a whole-brain dynamical model with 5HT2A-R neuromodulation. Furthermore, we built a simple model able to predict a region’s relative change in entropy from its local 5HT2A-R density and topological properties, showing that, somewhat paradoxically, at a whole-brain level receptor density is a poor predictor of 5HT2A-R activation effect.

Key to developing this predictive model was a three-way partition of brain regions according to their connectivity strength, suggesting a differentiated action mechanism of 5HT2A-R agonists that depends on the local topology of brain regions. In summary, our results suggest that the local changes in entropy, as well as the global entropy increase, induced by 5HT2A-R activation can be explained from a region-specific interplay between structural connectivity and receptor density. Finally, controlled experiments with null network models confirm that receptor density and connectivity strength are not only necessary, but also sufficient, to explain the entropic effects of 5HT2A-R activation.

The spatially heterogeneous, complex nature of the observed effects of 5HT2A-R activation opens a challenging problem for understanding the clinical and scientific relevance of psychedelic drugs, and implies a domain specificity to the so-called ‘entropic’ action of psychedelics.

## Methods

### Dynamic mean-field model with 5HT2A-R neuromodulation

The main computational tool used in this study is the Dynamic Mean-Field (DMF) model by Deco *et al.*,^18, 19^ which consists of a set of coupled differential equations modelling the average activity of one or more interacting brain regions. In this model, each brain region *n* is modelled as two reciprocally coupled neuronal masses, one excitatory and one inhibitory, and the excitatory and inhibitory synaptic currents *I*^(*E*)^ and *I*^(*I*)^ are mediated by NMDA and GABA_A_ receptors respectively. Different brain regions are coupled via their excitatory populations only, and the structural connectivity is given by the matrix *C*.

The full model, including the neuromodulatory effect described below, is given by

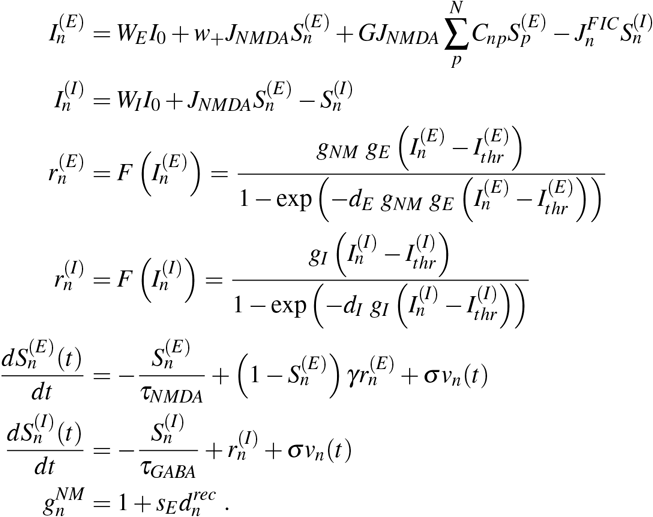

Above, for each excitatory (*E*) and inhibitory (*I*) neural mass, the quantities 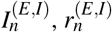, and 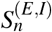 represent its total input current (nA), firing rate (Hz) and synaptic gating variable, respectively. The function *F* (·) is the transfer function (or *F-I curve*), representing the non-linear relationship between the input current and the output firing rate of a neural population. Finally, 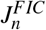 is the local feedback inhibitory control of region *n*, which is optimized^19^ to keep its average firing rate at approximately 3 Hz, and *v*_*n*_ is uncorrelated Gaussian noise injected to region *n*. The interested reader is referred to the original publication for further details.^19^

Key to this study is the recent extension of this model including neuromodulatory effects.^18^ In the equations above, 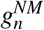 is a *neuromodulatory scaling factor* modulating the F-I curve of all brain regions in the model, which affects region *n* depending on its density of the receptor of interest, 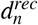. Including neuromodulation, the free parameters of the model are *G*, the global coupling parameter, and *s*_*E*_, the excitatory neuromodulatory gain, that are set to 2 and 0.2 respectively following Deco *et al*.^18^ The model was simulated using a standard Euler-Maruyama integration method,^40^ using the parameter values shown in Table 1.

**Table 1.** Dynamic Mean Field (DMF) model parameters

| *Parameter* | *Symbol* | *Value* |
| --- | --- | --- |
| External current | $I_0$ | 0.382 nA |
| Excitatory scaling factor for $I_0$ | $W_E$ | 1 |
| Inhibitory scaling factor for $I_0$ | $W_I$ | 0.7 |
| Local excitatory recurrence | $w_+$ | 1.4 |
| Excitatory synaptic coupling | $J_{NMDA}$ | 0.15 nA |
| Threshold for $F(I_n^{(E)})$ | $I_{thr}^{(E)}$ | 0.403 nA |
| Threshold for $F(I_n^{(I)})$ | $I_{thr}^{(I)}$ | 0.288 nA |
| Gain factor of $F(I_n^{(E)})$ | $g_E$ | 310 nC<sup>-1</sup> |
| Gain factor of $F(I_n^{(I)})$ | $g_I$ | 615 nC<sup>-1</sup> |
| Shape of $F(I_n^{(E)})$ around $I_{thr}^{(E)}$ | $d_E$ | 0.16 s |
| Shape of $F(I_n^{(I)})$ around $I_{thr}^{(I)}$ | $d_I$ | 0.087 s |
| Excitatory kinetic parameter | $\gamma$ | 0.641 |
| Amplitude of uncorrelated Gaussian noise $v_n$ | $\sigma$ | 0.01 nA |
| Time constant of NMDA | $\tau_{NMDA}$ | 100 ms |
| Time constant of GABA | $\tau_{GABA}$ | 10 ms |

### Parcellation, connectome, and 5HT2A receptor maps

In addition to the DMF equations above, to simulate whole-brain activity we need three more ingredients: a suitable parcellation of the cortex into well-defined regions, a structural connectivity matrix between those regions, and the density map of a receptor of interest - in our case, the serotonin 2A receptor.

As a basis for our simulation, we used the Automated Anatomical Labelling (AAL) atlas,^20^ a sulci-based parcellation of brain volume registered in MNI space. The AAL parcellation specifies a partition of the brain into 90 Regions Of Interest (ROIs), that provides sufficient level of detail to obtain a picture of the topographical distribution of entropy, while being suitable for computationally intensive simulations.

Structural connectivity data between the 90 AAL ROIs was obtained from 16 healthy subjects (5 female) using diffusion Magnetic Resonance Imaging (dMRI), registered to the MNI space and parcellated according to the AAL atlas on the subject’s native space. Then, for each subject the histogram of fiber directions at each voxel was obtained, yielding an estimate of the number of fibers passing trough each voxel. The weight of the connection between regions *i* and *j* was defined as the proportion of fibers passing through region *i* that reach region *j*, which (since the directionality of connections cannot be determined using dMRI) yields a symmetric 90×90 Structural Connectivity (SC) matrix. Finally, the SC matrix of each subject was thresholded, pruning any connection lower than 0.1%, and SC matrices of all subjects were averaged to obtain a single SC matrix used in all simulations. This average matrix was kindly provided by Deco *et al*.^18^

For the density map of 5HT2A receptors, we used the Positron Emission Tomography (PET) data made public by Beliveau *et al.*,^21^ which can be combined with the AAL parcellation to obtain an estimate of receptor density in every AAL region. Further details can be found in the original publications.^18, 21^

Together with the DMF equations, these three elements fully specify our whole-brain model and allow us to run simulations and obtain time series of excitatory firing rates, 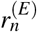, which we save for further analysis.

### Estimating the differential entropy of firing rates

The result of each simulation is a set of 90 time series representing the firing rate of each excitatory population 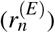, which we analysed to measure the entropy of the activity of each ROI. For a continuous random variable *X* with associated probability distribution *p*(*x*) and support 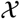, its differential entropy is defined as^41^

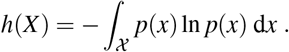

Estimating differential entropy from data is in general a hard problem, stemming (for the most part) from the difficulties in estimating a probability density from samples^42^. We found that the probability distributions of firing rates 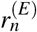 were well approximated by a gamma distribution: goodness-of-fit, measured with a standard Kolmogorov-Smirnov statistic, was satisfactory and comparable across brain regions and conditions (see Supp. Fig. 3).

This observation greatly simplifies the estimation of the differential entropy of each region, which is now reduced to estimating the shape and scale parameters of the gamma distribution, *k* and *θ*. Once these parameters are estimated (in our case by standard maximum likelihood estimation), the differential entropy of a gamma distribution can be computed analytically in closed form as

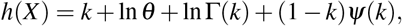

where Γ(*·*) and *ψ*(*·*) are the standard gamma and digamma functions, respectively.

### Linear models for Δ*h*_*n*_ and three-way separation of AAL regions

To explain the changes in differential entropy across regions and conditions, we trained several linear mixed-effects models^43^ using different sets of covariates designed to test specific hypotheses about the relation between entropy, 5HT2A-R density, and network topology.

First, we implemented a simple algorithm to find the group of regions where the change in entropy depends linearly on region strength. To do so, we designed an optimisation procedure to find the optimal linear fit between Δ*h*_*n*_ and connectivity strength after the removal of some regions from the fit by setting a threshold on 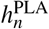. The procedure consisted in iteratively setting a threshold on 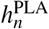; ii) fitting a linear model with strength as a regressor and Δ*h*_*n*_ as target for those regions below the threshold; and iii) computing the goodness of fit measured by *R*^2^. This process is repeated for the whole range of entropy values, yielding one *R*^2^ value per threshold. The threshold that maximises goodness of fit gives the optimal separation between regions, separating those regions whose change in entropy linearly depends on connectivity strength from those that it does not.

With this three-way grouping fixed, we investigated the power of several dynamical and topological features to explain the changes in local entropy (Δ*h*_*n*_) by including them as covariates in the linear model. The three-way separation of brain regions induces a model with 7 terms: a constant term and 2 fixed-effect terms for each group of regions: one for the connectivity strength and one for the receptor density. The model was fit by maximum likelihood using off-the-shelf software, and explanatory power measured using *R*^2^.

### Null network models of the human connectome

Null network models of the human structural connectivity were used to evaluate the role of the local connectivity properties^44^ on the local changes in entropy induced by 5HT2A-R activation. To this end, we applied three different randomisation schemes to the structural connectivity in order to produce suitable surrogate networks. These surrogates were designed to preserve different network attributes of the original connectome: i) the overall density and strength (RAND), ii) the degree distribution (degree-preserving randomisation [DPR]); and iii) the strength distribution (strength-preserving randomisation [DSPR]) After randomisation, the DMF model was run with and without 5HT2A-R activation, and entropies estimated and analysed following the same procedure as in the rest of the article. Every surrogate model was run 120 times, and the results averaged across runs.

In addition to the network surrogates, we computed several topological measures to include in the linear model reported in Fig. 3. These measures included betweenness, eigenvector centrality, closeness, communicability, PageRank index, and sub-graph centrality. All of these computations, including the surrogate networks and the topology measures, were performed using the Brain Connectivity Toolbox^45^.

## Acknowledgements

R.H. is funded by CONICYT scholarship CONICYT-PFCHA/Doctorado Nacional/2018-21180428. P.M. is funded by the Wellcome Trust (grant no. 210920/Z/18/Z). F.R. is supported by the Ad Astra Chandaria Foundation. R.C. is supported by CONICYT-PAI Inserción 79160120, Proyecto REDES ETAPA INICIAL, Convocatoria 2017 REDI170457 and Fondecyt Iniciación 2018 Proyecto 11181072.

## Author contributions statement

R.H. performed the numerical analysis and run the simulations. All the authors of this article conceptualized this research, design the methodology. All the authors carefully analysed the results, drafted and reviewed the manuscript.

