## Supplementary Figures for "A mechanistic model of the neural entropy increase elicited by psychedelic drugs"

### ABSTRACT

Psychedelic drugs, including lysergic acid diethylamide (LSD) and other agonists of the serotonin 2A receptor (5HT2A-R), induce drastic changes in subjective experience, and provide a unique opportunity to study the neurobiological basis of consciousness. One of the most notable neurophysiological signatures of psychedelics, increased entropy in neural activity, is thought to be of crucial importance to the psychedelic experience, mediating both acute alterations in consciousness and long-term effects on well-being – yet, no mechanistic explanation of this phenomenon has been put forward so far. In this paper we undertake this task, and build upon a recent whole-brain model of serotonergic neuromodulation to study the entropic effects of 5HT2A-R activation. Our results reproduce the overall entropy increase observed in previous experiments *in vivo*, providing the first mechanistic account of this phenomenon. We also found that entropy changes were not uniform across the brain: entropy increased in some regions and decreased in others, suggesting a topographical reconfiguration mediated by 5HT2A-R activation. Interestingly, at the whole-brain level this reconfiguration was not explained by 5HT2A-R density, but by topological properties of the brain's anatomical connectivity. These results help us understand the mechanisms underlying the psychedelic state and, more generally, the pharmacological modulation of whole-brain activity.

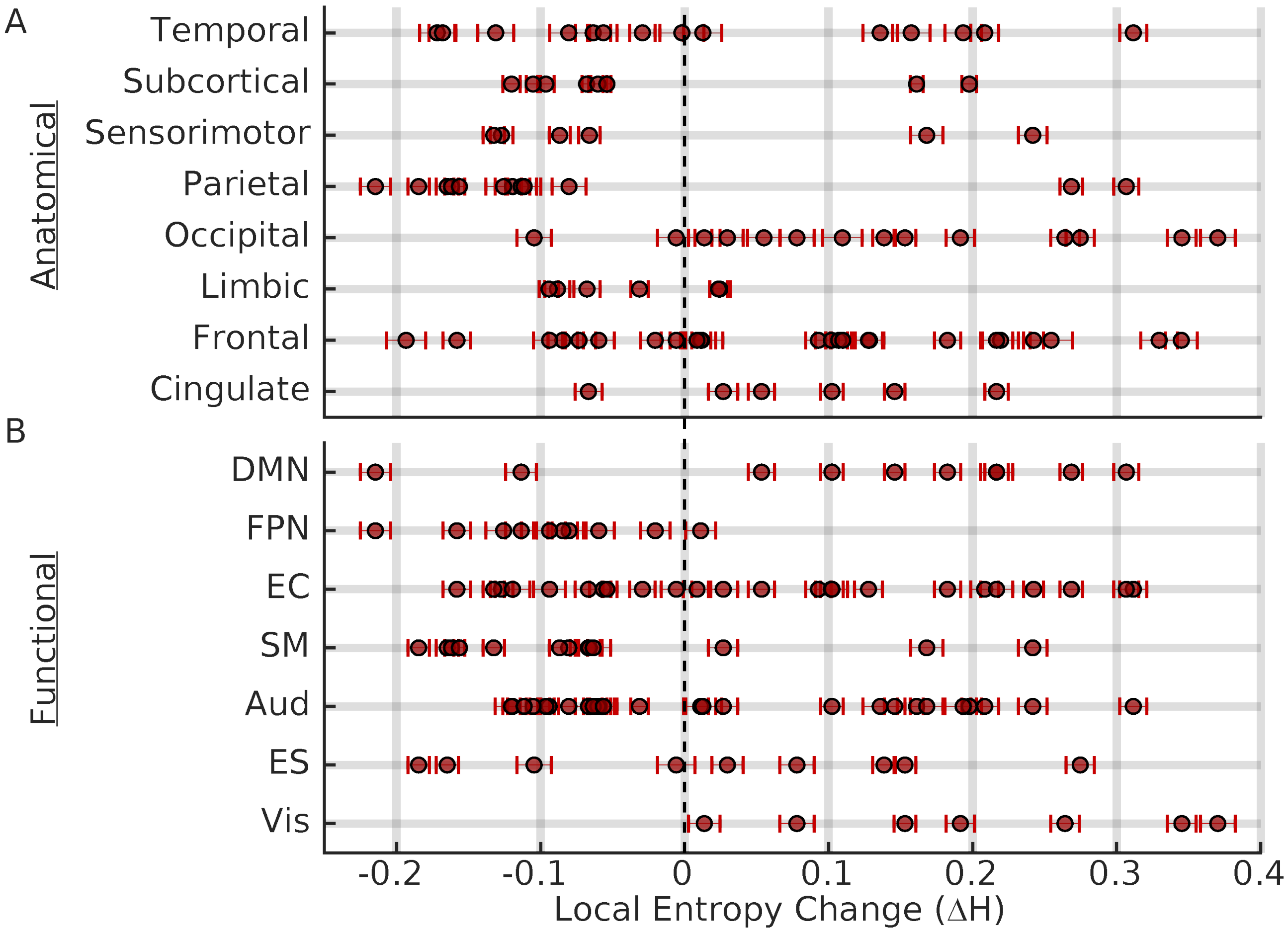


**Supplementary Figure 1. The effect of 5HT2A-R activation is not be explained by anatomical or functional groupings. A** 5HT2A-R activation heterogeneously change entropy in regions belonging to the same anatomical group. **B** Entropy also changed in a heterogeneous way when functional grouping is considered. Note the special case of FPN and Vis networks, for which almost all regions decreased and increased their entropy, respectively (note that functional groups can be overlapping). Circles are averages and error bars 1 s.d. out of 1000 simulations.

### Supplementary Figure 1. Heterogeneous entropy changes induced by 5HT2A-R activation on anatomical and functional groupings of brain regions.

We asked whether the heterogeneous changes of entropy induced by 5HT2A-R activation could be explained by grouping the AAL brain regions according to anatomical and functional criteria. Regarding anatomical criteria brain regions can be spatially split into 8 major non-overlapping anatomical groups: Frontal, Temporal, Parietal, Occipital, Limbic, Sensorimotor, Cingulate and Subcortical[1]. Regarding the functional grouping, we used the Resting State Networks[2], which is a functional grouping of brain regions based on the observed spatio-temporal patterns of BOLD signals during resting state activity. These groups are Fronto Parietal (FPN), Default Mode (DMN), Primary Visual (Vis), Extrastiate Cortex (EC), Auditory (Aud), Sensorimotor (SM), and Executive Control (EC). The first two, FPN and DMN, were obtained from Ref.[3] and the rest from Ref.[2]. Brain regions can potentially belong to different functional groups.

The effect of 5HT2A-R activation on entropy is heterogeneous also at the level of anatomical groups (Supplementary Figure 1A), i.e. within all the groups there were both regions with increased and decreased entropy after the 5HT2A-R activation. However, Occipital and Cingulate regions show a strong tendency to increase its entropy, in agreement with entropy increases in these regions observed in human experiments with serotonergic psychedelics[4,5]. Regarding the functional grouping (Supplementary Figure 1B), we also found that the effect of 5HT2A-R activation on regional entropy (Fig. 3B) is heterogeneous within groups, with the exception of Vis and FPN, where almost all the regions increased and decreased their entropy, respectively. Note also that with the exception of the Angular gyri, all the DMN regions increase their entropy, which resonates with the observed reduction of the DMN integrity on humans during psychedelic experiences[6].

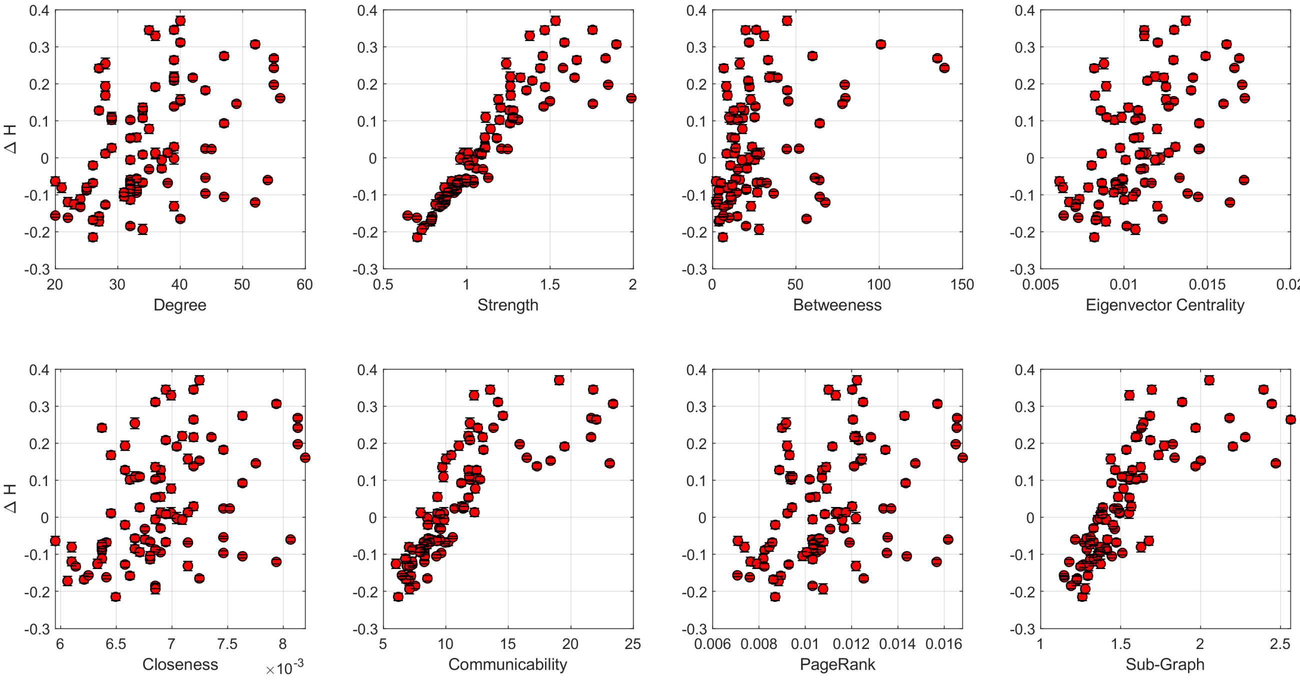


**Supplementary Figure 2. Centrality measures as possible explanatory variables for $\Delta H$.** Local connectivity strength is the best linear predictor of $\Delta H$ among other centrality measures. Communicability and sub-graph centrality also show a good linear relationship with $\Delta H$, but more spread, thus, they were not chosen. Circles are averages and error bars 1 s.d. out of 1000 independent simulations.

**Supplementary Figure 2. Connectivity strength is the best predictor for entropy changes among local connectivity measures.**

We control the role of local connectivity on the observed entropy changes ($\Delta H$) induced by 5HT2A-R activation using as predictors for $\Delta H$ other local connectivity measures as: degree[7], eigenvector centrality[7], communicability[8], page-rank[7], sub-graph centrality[7] and closeness centrality[9]. We confirmed that local connectivity strength is the best linear predictor of entropy changes among other centrality measures.

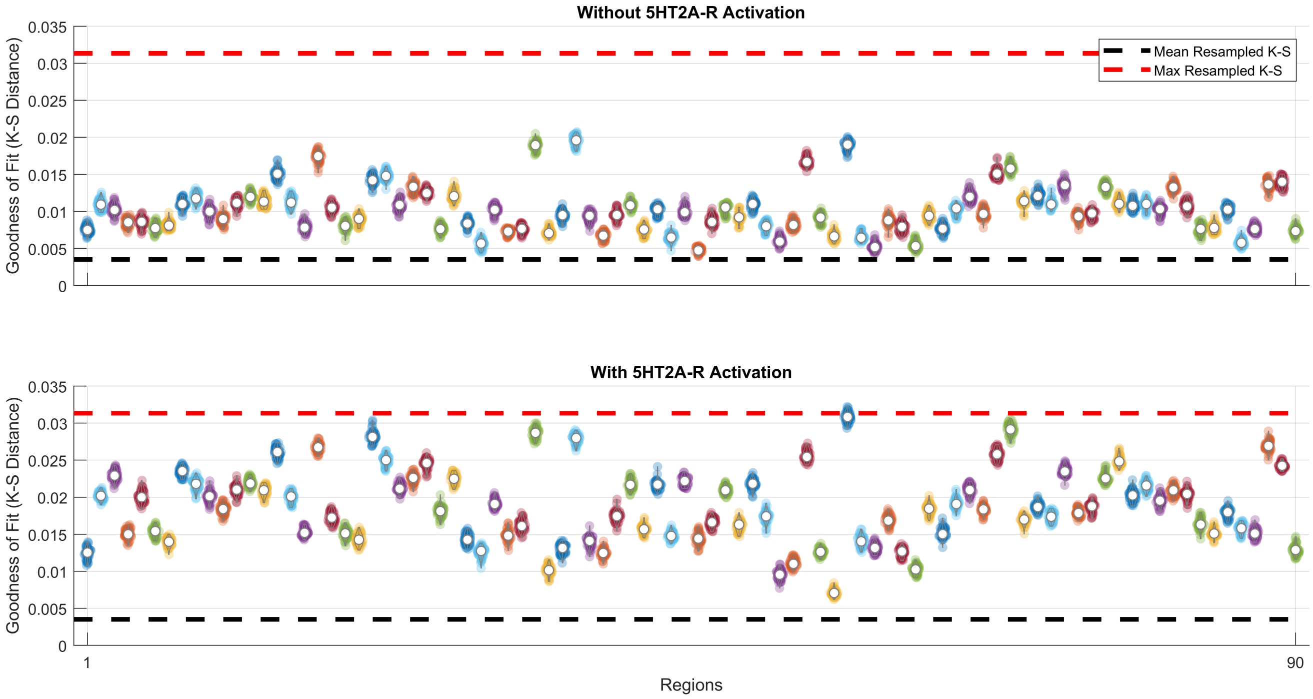


**Supplementary Figure 3. Goodness of fit of Gamma distribution to firing rates distribution are within confidence intervals.** Distribution of K-S distance values for PLA (top) and 5HT2A-R (bottom) condition for each region are represented by violins, where white circle denotes average K-S. Black (red) dashed line represent the average (maximum) re-sampled K-S distance (see the main text for details).

### Supplementary Figure 3. The probability distribution of excitatory firing rates can be well fitted by a Gamma distribution.

To check the goodness of fit (GOF) of Gamma distribution to the simulated excitatory firing rates of each region, we generated $10^6$ simulation points for each region under both PLA and 5HT2A condition, then, a Gamma distrbution was fitted and the Kolmogorov-Smirnov (K-S) distance between the firing rate distribution and 1000 random samples (same size) of the respective fitted Gamma distribution was computed. We summarize the Gamma GOF as the average K-S for each region under both conditions. To assess the significance of average GOF values, we generated a confidence interval computing the K-S distance between 100 independent samples (same size) sampled from exactly the same Gamma distribution (re-sampled K-S). This procedure was repeated for different set of Gamma distribution parameters. If the average K-S distance of a given region falls below the maximum re-sampled K-S distance, we consider this region to be well fitted by the Gamma distribution.

Despite the decreased GOF under 5HT2A condition (compared to PLA) all regions fall within the confidence interval under both conditions, enabling us to use the Gamma distribution parameters to estimate each region's Shannon's differential entropy.
